# EukProt: A database of genome-scale predicted proteins across the diversity of eukaryotes

**DOI:** 10.1101/2020.06.30.180687

**Authors:** Daniel J. Richter, Cédric Berney, Jürgen F. H. Strassert, Yu-Ping Poh, Emily K. Herman, Sergio A. Muñoz-Gómez, Jeremy G. Wideman, Fabien Burki, Colomban de Vargas

## Abstract

EukProt is a database of published and publicly available predicted protein sets selected to represent the breadth of eukaryotic diversity, currently including 993 species from all major supergroups as well as orphan taxa. The goal of the database is to provide a single, convenient resource for gene-based research across the spectrum of eukaryotic life, such as phylogenomics and gene family evolution. Each species is placed within the UniEuk taxonomic framework in order to facilitate downstream analyses, and each data set is associated with a unique, persistent identifier to facilitate comparison and replication among analyses. The database is regularly updated, and all versions will be permanently stored and made available via FigShare. The current version has a number of updates, notably ‘The Comparative Set’ (TCS), a reduced taxonomic set with high estimated completeness while maintaining a substantial phylogenetic breadth, which comprises 196 predicted proteomes. A BLAST web server and graphical displays of data set completeness are available at http://evocellbio.com/eukprot/. We invite the community to provide suggestions for new data sets and new annotation features to be included in subsequent versions, with the goal of building a collaborative resource that will promote research to understand eukaryotic diversity and diversification.

## Introduction

Over the past 15 years, the discovery of diverse novel microbial eukaryotes, coupled with methods to reconstruct phylogenies based on hundreds of protein-coding genes (frequently referred to as phylogenomics, although the term was originally proposed as the more general integration of genome analysis and evolutionary studies [Eisen, 2003]) have led to a remarkable reshaping in our understanding of the eukaryotic tree of life, the proposal of new supergroups and the placement of enigmatic lineages in known supergroups (Burki et al., 2012, 2020; Kamikawa et al., 2014; Yabuki et al., 2015; Janouškovec et al., 2017; Brown et al., 2018; Lax et al., 2018; Strassert et al., 2019; Gawryluk et al., 2019; Tice et al., 2021). The phylogenomic approach has also been used to investigate branching patterns within eukaryotic supergroups, including to understand species evolutionary relationships with implications for the early evolutionary events in the three best-studied eukaryotic lineages: land plants (Wickett et al., 2014), animals (King & Rokas, 2017) and fungi (Kiss et al., 2019; Li et al., 2021; Strassert & Monaghan, 2022). Furthermore, analyses in diverse eukaryotes have the potential to reveal new genes and pathways for biological processes currently characterized only in these three well-studied lineages and their parasites (del Campo et al., 2014; Richter & Levin, 2019). In the ocean alone, planetary-scale metagenomics studies (Bork et al., 2015; Tara Oceans Coordinators et al., 2020) have already unveiled an extreme diversity of eukaryotic genes (Carradec et al., 2018) whose phylogenetic origin and ecological function are mostly unknown.

A critical prerequisite to all genomic studies is the underlying database of predicted proteins from which orthologs are extracted or other sequence analyses are performed. Because no single source contains the protein predictions for the complete set of eukaryotes that have been sequenced at a genomic scale, each study needs to assemble their own database, reducing reproducibility among analyses (due to the inclusion of different taxa, or different data sets for the same taxon), and producing a significant barrier to new researchers entering the field. In addition, because the large majority of the protein data sets from diverse eukaryotes are not included in major public databases (see Figure 2), researchers cannot easily access them via standard tools such as NCBI BLAST (Sayers et al., 2020) in order to find the homologs of a protein of interest, nor are they automatically integrated into major public databases containing annotations such as protein domains (e.g., Pfam [El-Gebali et al., 2019], Interpro [Mitchell et al., 2019]) or gene ontology (The Gene Ontology Consortium, 2019).

To address this gap, we assembled a database of protein sequences – EukProt – either from newly performed protein prediction or by retrieving available sequences from a comprehensive set of species representing known eukaryotic diversity. We note that an existing database of genome-scale protein data sets, PhyloDB (https://github.com/allenlab/PhyloDB), contains 550 eukaryotic species with at least 500 proteins, as of version 1.076. PhyloDB was most recently updated in 2015; EukProt includes data made available since then, allowing it to provide more species with generally higher completeness and representing a greater phylogenetic breadth. In addition, we placed each species within a universal eukaryotic taxonomic framework, UniEuk (Berney et al., 2017), in order to ensure that the evolutionary relationships among data sets are accurately and consistently described. We also provide estimated completeness statistics for each predicted proteome based on BUSCO scores and have selected a suggested subset of 196 proteomes for use in comparative genomics investigations. EukProt is designed to prioritize ease of use, with unique, persistent identifiers assigned to each data set and a standard system of nomenclature to facilitate repeatability of analyses. To this end, we have followed FAIR principles (Findable, Accessible, Interoperable, Reusable) in its construction (Wilkinson et al., 2016).

## Methods

### Species and strain identity

We determined species and strain identities by reading the publications that described the data sets, consulting the literature for naming revisions, and comparing 18S ribosomal DNA sequences for each data set to reference sequence databases (see below for a description of how we retrieved 18S sequences). For species that were previously known by other names, we recorded these previous names in the metadata for the data set, except in cases where a species was originally assigned to a genus but not identified to the species level (e.g., *Goniomonas* sp., now identified as *Goniomonas avonlea*, is not listed as a previous name). This exception was not followed when the GenBank record description for the 18S sequence contained a different name than the one used in EukProt; in this case this previous name is always listed in the previous names field, in order to avoid confusion when retrieving 18S sequences from GenBank.

When no strain name was available for an MMETSP data set, we used the MMETSP ID as the strain name (for example, EP00362 has the strain name MMETSP1317). For strains with alternative names, we included these in a dedicated field for alternative strain names; this list is not necessarily exhaustive (i.e., it is not guaranteed to contain all alternative names for a given strain).

### Supergroups and taxogroups from UniEuk

The full taxonomic pathways provided for all species (which follow the framework developed in the UniEuk project [Berney et al., 2017]) are not based on a fixed number of ranks, but on a free, unlimited number of taxonomic levels, in order to match phylogenetic evidence as closely as possible. This provides end-users more information and flexibility, but could also make it more difficult to summarize results of downstream analyses. Therefore, we provide three additional fields (“supergroup”, “taxogroup1” and “taxogroup2”) to help end-users whenever it is useful to distribute eukaryotic diversity into a fixed number of taxonomic categories of roughly equivalent phylogenetic depth or ecological relevance. The groupings called “supergroups’’ in the context of UniEuk resources (36 recognized lineages so far, of which 34 are included in EukProt; *Microheliella* and *Meteora* are not represented) consist of strictly monophyletic, deep-branching eukaryotic lineages of a phylogenetic depth equivalent to some of the first proposed supergroups such as Opisthokonta, Alveolata, Rhizaria, and Stramenopiles. UniEuk “supergroups” correspond roughly to the deepest phylogenetic resolution of eukaryotic relationships achievable with ribosomal RNA genes, and not necessarily to the currently recognized highest-level groupings of eukaryotes based on phylogenomic evidence (these groupings are, of course, present in the complete taxonomic pathways). UniEuk “supergroups” are therefore highly variable in relative diversity, ranging from lineages consisting of a single, orphan genus (e.g., *Ancoracysta, Mantamonas, Palpitomonas*), to Opisthokonta as a whole. The “taxogroup1” (of which there are 72 in EukProt) and “taxogroup2” (of which there are 198 in EukProt) levels allow further subdivision of large supergroups into lineages of relatively equivalent evolutionary or ecological relevance, based on current knowledge. These levels are more arbitrarily defined but are intended to represent strictly monophyletic groupings that match one of the levels in the complete taxonomic pathways. As an illustrative example, diatoms are in the Diatomeae taxogroup2, which is within the Ochrophyta taxogroup1, which is within the Stramenopiles supergroup. Small, ecologically and morphologically homogeneous supergroups are not subdivided further; in such cases the “taxogroup1” and “taxogroup2” levels are the same as the “supergroup” level. The same approach will be used in EukRibo, a manually-curated database of reference ribosomal RNA gene sequences developed in parallel (Berney 2022) to help users link analyses of different types of genetic data.

### Merging strains from the same species

In general, we only included data from a single strain/isolate per species. However, when only a single transcriptome data set was available for a given strain of a species, and there were additional published transcriptome data sets for other strains of the same species, we combined them using CD-HIT (Li & Godzik, 2006) run with default parameter values, in order to guard against the possibility that a single transcriptome might lack genes expressed only in one condition or experiment. When multiple strains were merged to produce a species’ data set (there were 41 such cases), this information is indicated in the metadata for the data set. When a data set is indicated as merged, but more than one strain name is not listed in the strain column, this indicates that the data set was distributed as merged and we were unable to determine the exact strains used.

### Processing steps applied to publicly available data

All sequences within each FASTA file are assigned a unique, standardized identifier based on the data set’s EukProt ID and on the type of data (protein or transcriptome); this identifier is prepended to the existing FASTA header, separated by a space. Illegal characters are removed from sequences. The following characters are permitted, as defined by NCBI BLAST (https://blast.ncbi.nlm.nih.gov/Blast.cgi?CMD=Web&PAGE_TYPE=BlastDocs&DOC_TYPE=BlastHelp) for nucleic acid sequences: ACGTNUKSYMWRBDHV and for protein sequences: ABCDEFGHIKLMNPQRSTUVWYZX*.

EukProt metadata indicate the additional steps applied to each data set after downloading from the data source (if any).

All software parameter values were default (unless otherwise specified below or in the metadata record for a given data set), as default parameter values are most likely to correspond to the options recommended for general use by the authors of the software. Due to the large volume of data sets we processed, and variability among them, we were not able to test parameter values beyond those specified below.

#### assemble mRNA

*De novo* transcriptome assembly using Trinity v. 2.8.4, http://trinityrnaseq.github.io/ (Haas et al., 2013). We trimmed Illumina input reads for adapters and sequence quality using the built-in ‘--trimmomatic’ option (whose default trimming settings are based on optimal trimming parameters from [MacManes, 2014]). We trimmed 454 input reads prior to running Trinity with Trimmomatic v. 0.3.9, http://www.usadellab.org/cms/?page=trimmomatic (Bolger et al., 2014) with the directives ‘ILLUMINACLIP:[454 adapters FASTA file]:2:30:10 SLIDINGWINDOW:4:5 LEADING:5 TRAILING:5 MINLEN:25’ (corresponding to the default Trinity trimming parameters, but for single-end reads). When the sequence library was described as stranded in the NCBI Sequence Read Archive, we used the corresponding ‘SS_lib_type’ option.

#### translate mRNA

*De novo* translation of mRNA sequences with Transdecoder v. 5.3.0, http://transdecoder.github.io/. When the number of predicted protein sequences for a given species was both less than half of the input mRNA sequences and less than 15,000, we reduced the minimum predicted protein length to 50 (from the default of 100).

#### CD-HIT

Clustering of protein sequences to produce a non-redundant data set using CD-HIT v. 4.6, http://weizhongli-lab.org/cd-hit/ (Li & Godzik, 2006). We used this tool principally to combine protein predictions for different strains of the same species, but also to reduce the size of very large predicted protein sets (>50,000 proteins) that showed evidence of redundancy.

#### extractfeat, seqret, transeq, and trimseq

From the EMBOSS package v. 6.6.0.0, http://emboss.sourceforge.net/ (Rice et al., 2000). We used extractfeat to produce coding sequences (CDS) from genomes with gene annotations in EMBL format but without publicly available protein sequences. We used seqret to convert FASTQ files to FASTA files. We used transeq to translate CDS directly into proteins. We used trimseq to trim EST sequences before translation with Transdecoder.

#### gffread

To produce protein sequences from genomes with gene annotations in GFF format but without publicly available protein sequences, using the command line option ‘-y’ in version 0.12.3 (Pertea & Pertea, 2020).

#### predict genes

We used EukMetaSanity https://github.com/cjneely10/EukMetaSanity (Neely et al., 2021) to perform automated annotation of genome sequences lacking publicly available protein predictions. We used the following parameters, as specified in (Alexander et al., 2021): --min_contig 500 --min_contig_in_predict 500 --max_contig 100000000. These settings were designed for generalized gene prediction on genomes expected to be sampled from across eukaryotic diversity, which is also the case for the data sets in EukProt. We used the parameter --min_contig_in_predict 200 (as it matched the default minimum contig length in Trinity). By default, we selected the proteins at Tier 2 (predictions supported by at least 2 sources). If Tier 2 produced fewer than 15,000 predicted proteins, we instead selected Tier 1. All other parameter values were left at their defaults. We did not perform gene prediction on unannotated genomes for which a transcriptome was already available for the same species (under the assumption that the gene predictions of the transcriptome would be of higher quality, due to potential errors in the gene annotation process).

### Retrieval of 18S sequences

We searched for 18S sequences of at least 1,500 base pairs in length for each data set in the following order. If no sequence, or only a partial sequence was retrieved in a given step, we moved on to the next step. Details specific to each data set are included in the EukProt metadata.

1. Search GenBank (Sayers et al., 2020) for an 18S sequence for the species and strain. When more than one sequence was available for the strain, we chose the one containing the largest quantity of high-quality nucleotides (after identifying and eliminating sequencing errors, chimeras, etc.).
2. If the data set was sequenced as part of the MMETSP, use the 18S sequence from MMETSP (if it was provided in the project metadata, and also was not already present in GenBank).
3. Search GenBank for an 18S sequence for another strain of the same species.
4. Using the 18S sequence for the most closely related species in GenBank as a query, search the contigs associated with the data set (either genome or transcriptome) using blastn (Altschul, 1997).
5. Assemble the 18S sequence from the raw sequence reads associated with the data set, using phyloFlash version 3.4 (Gruber-Vodicka et al., 2020) with the SILVA database version 138.1 (Quast et al., 2012) and SPAdes assembler version 3.14.1 (Prjibelski et al., 2020), with default parameter values except for the number of reads used, and with read length set to match the read length of the data set. For each data set, we initially ran phyloFlash with all sequence reads. If we observed that there was evidence for reads mapping to the 18S of the target lineage, but that did not result in a contig assembled by SPAdes, we ran phyloFlash again with only the first 10 million reads. If this run showed a similar result (reads mapping but no contigs), we ran again with only the first 1 million reads. When all three runs (all reads, first 10 million reads, first 1 million reads) did not result in a SPAdes assembly, we selected the reads mapping to the 18S of the target lineage and performed an assembly using CAP3 version date 02/10/15 (Huang, 1999) with ‘relaxed’ parameter values from (Close et al., 2009): -p 75 -d 200 -f 250 -h 90.

### A selected subset of EukProt for phylogenetic studies

Single-copy orthologs were identified in EukProt v3 proteomes using BUSCO v5.2.2 (Manni et al., 2021) with the Eukaryota Odb10 database. BUSCO searches were run in protein mode with the default E-value cutoff of 1e-3. These BUSCO scores were visualized using Plotly (Plotly Technologies Inc., 2015) and are available at http://evocellbio.com/eukprot/. Single-copy BUSCO markers for taxa with BUSCO scores >30 were aligned to single-copy BUSCO profile HMMs with hmmalign (HMMER v3.3; [Eddy, 2020]), trimmed with ClipKit (-m gappy; [Steenwyk et al., 2020]), and concatenated into a supermatrix with FASConCat-G v1.04 (Kück & Longo, 2014). A guide tree was first inferred from this supermatrix with IQ-TREE v2.0.3 using the LG4X model (-fast mode; [Minh et al., 2020]). A second phylogenetic tree was inferred using the LG+PMSF(C10)+F+G4 model and using the LG4X tree as a guide (Wang et al., 2018). A first subset of taxa was automatically selected from the previously inferred tree to maximize phylogenetic diversity using Treemmer v0.1 (Menardo et al., 2018). A few taxa that were phylogenetically misplaced with high support, likely due to a high percentage of contaminating sequences, were visually detected and manually removed. Furthermore, predicted proteins from model organisms (e.g., *Homo sapiens, Saccharomyces cerevisiae, Toxoplasma gondii, Dictyostelium discoideum*, etc.) were included in the TCS. In addition, phylogenetically important taxa with sufficiently high BUSCO scores were also included. Since different lineages universally lack particular BUSCOs, a singular score cut-off is not reasonable and therefore each taxon group was assessed in relation to a model system with a trusted genome. For example, while *Giardia intestinalis* and *Toxoplasma gondii* have low BUSCO scores (23.5 and 61.2, respectively), they are generally accepted as complete genomes. Thus, species in these lineages (e.g., Fornicata, Preaxostyla, Apicomplexa, Parabasalia, etc.) with comparable BUSCO scores were included in the TCS. Finally, in taxonomically important lineages in which no model yet exists (e.g., Ancyromonads) proteome sets with BUSCO scores ~50% or greater were included, or if no such representative was available, it was excluded from the TCS (e.g., Tsukubomonads). We welcome suggestions from the community for future iterations of the TCS.

### The EukProt database distribution on FigShare

The database is distributed in five files. One file contains 993 protein data sets, for species with either a genome (375), a single-cell genome (56), a transcriptome (498), a single-cell transcriptome (47), or an EST assembly (17). A second file contains assembled transcriptome contigs, for 126 species with publicly available mRNA sequence reads but no publicly available assembly. The proteins predicted from these assemblies are included in the proteins file. A third file contains GFF annotations for 40 genomes lacking publicly available predicted protein annotations. Finally, the database metadata are distributed as two files: one file for the data sets included in the current version of the database (993), and a second file for data sets not included (163), accompanied by the reason they were not included. For example, when a data set cleaned with a genome-scale decontamination process is described in a manuscript, but the decontaminated data are not publicly available, it is not included in the database.

## Results and Discussion

### The EukProt database

EukProt (currently in version 3; 22 November, 2021) contains 993 eukaryotic species from 5 different sequence data-types (Figure 1): genome (375 species), single-cell genome (56), transcriptome (498), single-cell transcriptome (47), and expressed sequence tag (EST; 17). The data sets were downloaded from 37 different sources (Figure 2), with the two principal sources being NCBI (Sayers et al., 2020) and the Marine Microbial Eukaryote Transcriptome Sequencing Project (MMETSP) (Keeling et al., 2014). The database contains broad taxonomic representation, with species from 34 of 36 major deeply-branching eukaryotic lineages defined in the UniEuk database (Berney et al., 2017). Within these major groups, however, the representation remains uneven, which results from several factors: the difficulty of discovering, culturing or sequencing species from many lineages; a bias towards sequencing either macroscopic, multicellular species or unicellular species that are parasites (e.g., Apicomplexa), photosynthetic (such as diatoms, which are part of the ochrophyte lineage), or are otherwise economically important (del Campo et al., 2014); and very uneven evolutionary histories among groups, with some having undergone major diversification with many different morphologies, and others having diversified to a lesser extent (or with only a few surviving lineages).

**Figure 1:**
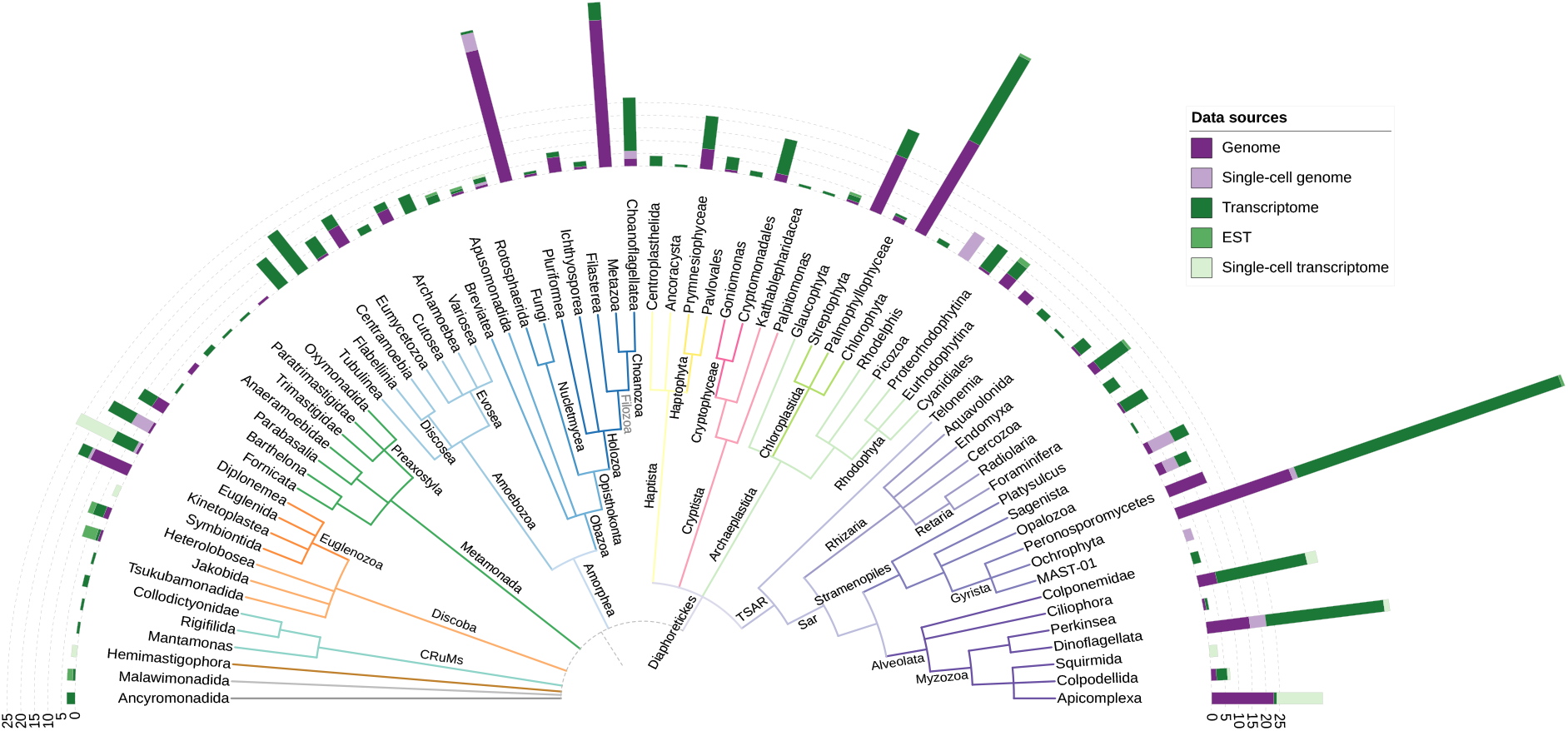
distribution of 993 source data sets on the eukaryotic tree of life, separated by data set type, with taxonomy based on UniEuk (Berney et al., 2017; Adl et al., 2019). The position of the eukaryotic root, indicated with a dashed line, is currently unresolved. Group names shown in gray are not officially recognized. The figure was created with iTOL (Letunic & Bork, 2019).

**Figure 2:**
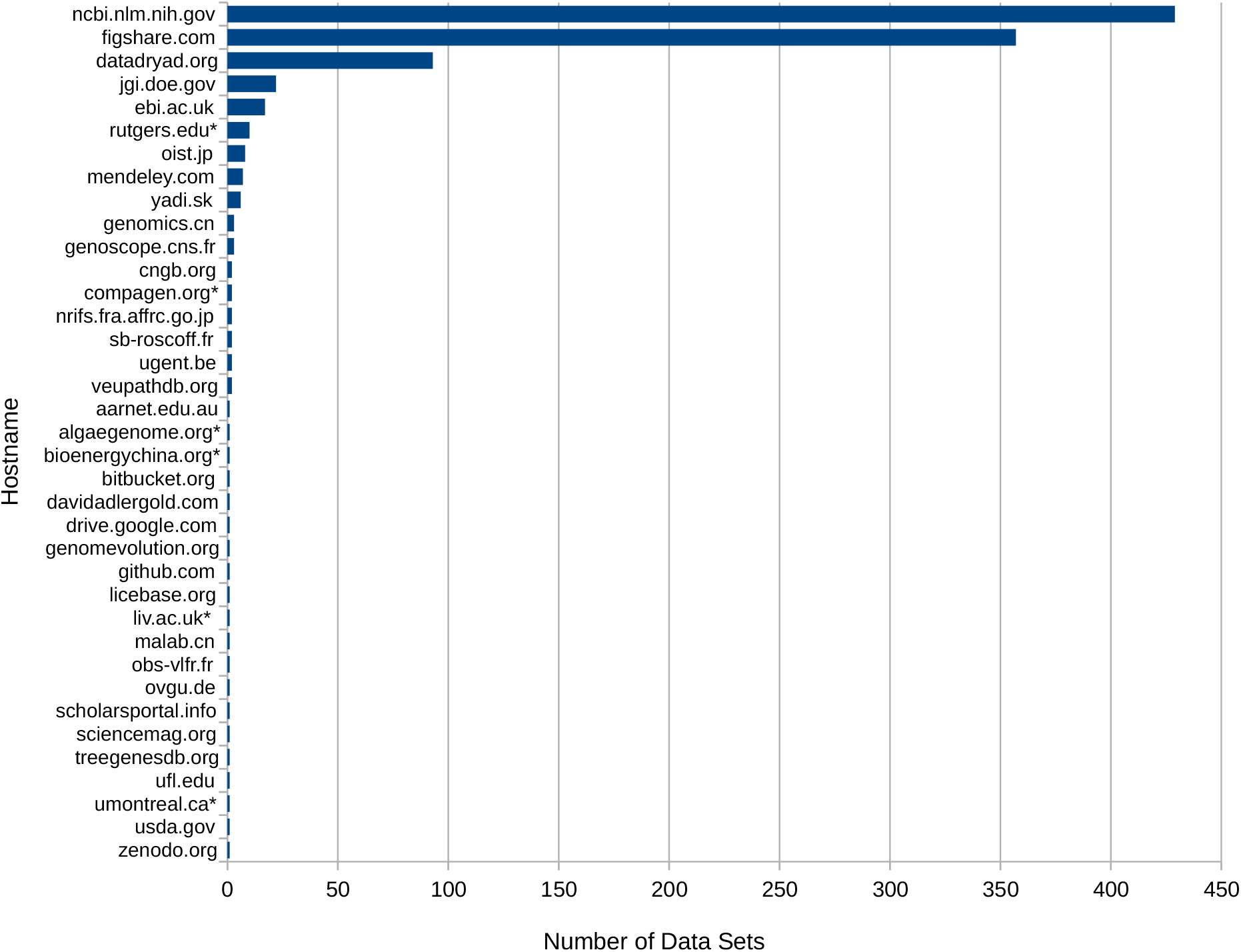
the 37 different web hosts from which data sets were downloaded. The count for figshare.com includes 272 species from the MMETSP (Keeling et al., 2014) for which a procedure to remove cross-contamination was applied (Marron et al., 2016). Individual URLs for each data set are listed in the metadata files available on FigShare. * indicates that the URLs for one or more data sets point to servers that are no longer responding or are otherwise inaccessible.

We have implemented in version 3 several changes proposed by the community of EukProt database users after version 2 was released: we have included (i) the 18S ribosomal DNA sequence corresponding to each data set (when available), (ii) a measurement of the completeness of the data set as estimated by BUSCO (Simão et al., 2015; Manni et al., 2021), and (iii) the assignment of unique EukProt identifiers to individual protein sequences.

The EukProt database is organized around 5 guiding principles:

#### Breadth of species phylogenetic diversity

The objective of the database is to include as many lineages as possible from the broad diversity of eukaryotes. For the majority of lineages, at least one representative from all sequenced genera is included. The exceptions are lineages with hundreds of sequenced representatives, or lineages whose sequenced species are relatively closely related, for which we included only a subset of species selected to represent phylogenetic diversity in order to balance the overall size of the database versus broad representation of eukaryotes. These well-sequenced lineages are Chloroplastida (containing the land plants and green algae), Metazoa (animals), Amoebozoa, Fungi, and some parasites/pathogens (members of Apicomplexa, Peronosporomycetes, Metakinetoplastina and Fornicata). Within animals, we also emphasized the inclusion of marine planktonic species, as we anticipate these could serve as mapping targets for data from large-scale metagenomic and metatranscriptomic ocean sequencing projects (e.g., *Tara* Oceans [Karsenti et al., 2011; Carradec et al., 2018; Richter et al., 2019], Malaspinas [Duarte, 2015; Acinas et al., 2021], Bio-GO-SHIP [Larkin et al., 2021] and GEOTRACES [Biller et al., 2018]).

#### Convenience of access

The full data set and its associated metadata can easily be downloaded from FigShare (Richter, Berney et al., 2022a). For 624 of the total 993 species in the database, publicly available protein sequences were ready to be included directly. For the remaining 369 species, additional processing steps were required to produce protein sequences for inclusion in the database. The most common bioinformatics processing steps were *de novo* assembling and translating transcriptomes from raw read data (120), translating mRNA sequences from publicly available assembled transcriptomes or EST projects lacking protein predictions (102), merging independent sets of protein predictions for the same species (85 species) and, newly added in version 3 of the database, producing *de novo* gene predictions for unannotated genomes (40). We also provide transcriptome assemblies and GFF annotation files for the latter two classes, as they are not publicly available. A full list of the different types of actions is described in the Methods, and the actions taken for each species are available in the database metadata.

#### A unified taxonomic framework

We placed all species into the taxonomic framework generated in the UniEuk project (Berney et al., 2017), which is based on the most recent consensus eukaryotic classification (Adl et al., 2019) and phylogenomic evidence. This will maximize interoperability of the database with other resources built upon to the UniEuk taxonomic framework (e.g., EukBank and EukMap [Berney et al., 2017], and EukRibo [Berney, 2022]), and ensure that results arising from EukProt can use a common set of identifiers to describe analyses at different taxonomic levels. For example, this might include labeling groups in a phylogenetic tree, or summarizing read placement data by taxonomic group. Knowledge of the taxonomy can also facilitate the detection of mis-identified gene sequences within a data set, via comparison of the topology of gene trees to the taxonomy. To assist in species identification and to provide a convenient link to EukRibo (a manually curated database of 18S rDNA sequences, part of UniEuk, aimed at taxonomic annotation of metabarcoding data sets), we also supply in version 3 the 18S sequence corresponding to the sequenced strain (or, if an 18S for the sequenced strain is not available, an 18S sequence from the same species; see Methods). In addition to the complete taxonomic pathway containing all intermediate nodes, we provide for each species three sequentially nested groupings (“taxogroup2” within “taxogroup1” within “supergroup”; see Methods) that can help to judge the redundancy of data sets, or conversely their lack of representation, within different groupings. The taxogroup1 level is used as the leaves in Figure 1. Finally, for each data set we include a list of previous species identifications, if any, to prevent issues resulting from revisions to species names or lineages and corrections of species misidentifications, which have occurred frequently with improvements in sequencing and phylogenetic techniques and the discovery of new eukaryotic organisms.

#### Appropriate references to publications and data sources

To ensure that the researchers who generated and provided each data set receive appropriate credit, we provide the DOI of the publication describing each data set as well as the URL from which it was downloaded. The list of URLs should also allow users of the database to download the original sequences for each data set (although it is not possible to guarantee that the URLs for all data providers will be permanently available).

#### Reusability, persistence and replicability

The database will be released in successive versions, each of which will be permanently stored and accessible at FigShare (Richter, Berney et al., 2022a). Thus, analyses using the database will need only to specify which version was used, enabling follow-up analyses or replications to begin with the identical database. In addition, each individual data set within the database is assigned a unique, permanent identifier. When a new data set becomes available for a given species, it is assigned a new unique identifier (and the identifier of the data set it replaces, if any, is indicated in the database metadata). These and all other changes between versions of the database are recorded as appropriate.

### ‘The Comparative Set’ (TCS): A selected subset of EukProt for comparative genomics investigations

Comparative genomics investigations are crucial to bridge the gaps between cell biology and biochemistry on one hand and diversity and evolution on the other. Classically, the set of species to include has been chosen by the authors of each study. With the goal of providing some consistency in future comparative genomics studies, we have selected a set of predicted proteomes with high completeness to represent the broad phylogeny of eukaryotes. A total of 196 predicted proteomes were selected as ‘The Comparative Set’ (TCS; Figure 3), abbreviated in honor of the late Tom Cavalier-Smith (Saldarriaga, 2021; Bass, 2021; Richards, 2021; Roger, 2021). All major model systems were included in this set as well as proteomes derived from high-quality genomes. Additional proteomes were chosen to maximize phylogenetic diversity and based on their completeness as estimated by BUSCO (Manni et al., 2021) scores. Updates to the TCS will be based on availability of new proteomes from phylogenetically important lineages currently underrepresented, and on community feedback. Please send any comments or suggestions for the TCS to. The complete EukProt database and the TCS are available for download and also available for BLAST searches at http://evocellbio.com/eukprot/ using Sequenceserver (Priyam et al., 2019).

**Figure 3:**
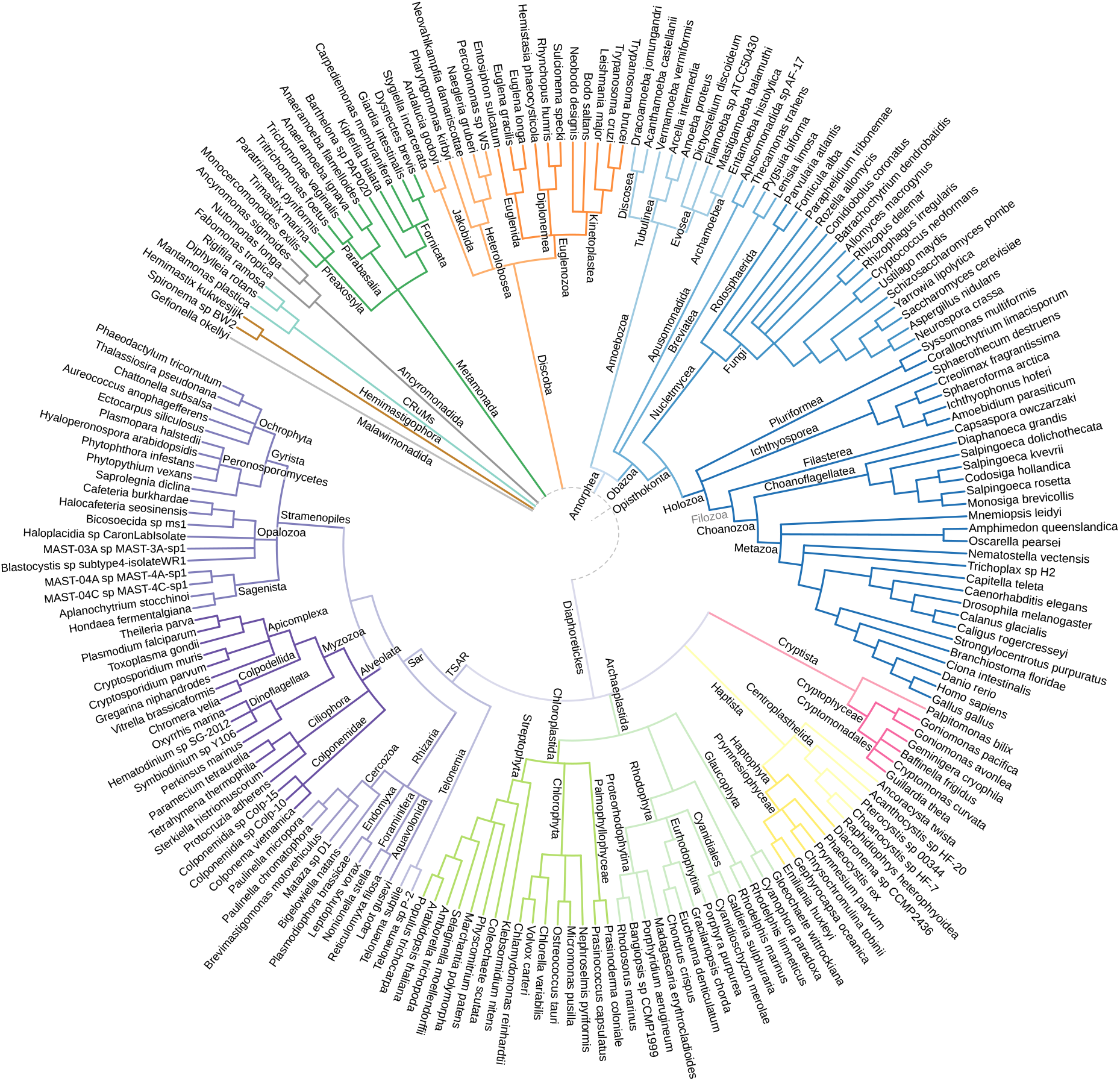
distribution of 196 selected data sets of The Comparative Set (TCS) on the eukaryotic tree of life, with relationships based on the UniEuk taxonomy (Berney et al., 2017; Adl et al., 2019). The position of the eukaryotic root, indicated with a dashed line, is currently unresolved. Group names shown in gray are not officially recognized. The figure was created with iTOL (Letunic & Bork, 2019).

### Current limitations of the EukProt database

In the course of constructing successive versions of EukProt, we have observed several areas that we believe to be current limitations of the database:

1. As can be observed in Figure 1, representation of species is highly uneven among taxonomic groups. This is largely due to the difficulty in identifying, cultivating and sequencing species from underrepresented groups; however, some representatives of key taxa have been published but are not publicly accessible (a list of these species can be found in the “not included” metadata table of our FigShare distribution).
2. We currently lack an automated, consistent method to identify and remove contaminated data sets or individual contaminant sequences. Currently, contaminated data sets are identified by end users and removed in subsequent versions.
3. Most species are not represented by genomes, which are the gold standard for providing gene catalogs. Instead, due to technical limitations in cultivation and/or sequencing methods, they are represented by transcriptomes from laboratory cultures or by single-cell genomes or transcriptomes.
4. Proteins smaller than 50 amino acids are not predicted with the settings we used for protein prediction (but may be included in data sets for which we did not perform protein prediction). We used the default settings for protein prediction in TransDecoder (see Methods), in order to compromise between minimum protein length and total number of predicted proteins (as proteomes predicted with protein sizes smaller than 50 amino acids are generally much larger, which may significantly slow many downstream analyses). EukProt users interested in proteins shorter than 50 amino acids (for the species on which we performed protein predictions) would instead have to repeat protein predictions using their desired settings.

### Growing the EukProt database with community involvement

The core functionality of the database is the distribution of genome-scale protein sequences across the diversity of eukaryotic life and within the UniEuk framework. However, we anticipate that numerous other features might be useful to the community, and we hope to involve the community in suggesting and adding new features in successive versions. These may include information or analyses on full data sets, such as the sequencing technology that was used (e.g., Illumina, PacBio), or the estimation of potential contamination levels (e.g., from mixed cultures or prey species) with non-target species, as inferred using systematic sequence homology searches. Additionally, we propose to use our GitHub (Richter, Berney et al., 2022b) as a flexible repository to disseminate community-contributed annotations on the level of individual protein sequences, such as protein domains from Pfam (El-Gebali et al., 2019)/Interpro (Mitchell et al., 2019), gene ontology (The Gene Ontology Consortium, 2019)/eggNOG (Huerta-Cepas et al., 2019), which may be updated in between EukProt releases.

As new genome-scale eukaryotic protein data sets become available, we plan to add them to the database. As yet, we do not have a formal mechanism to accomplish this, and will instead depend on monitoring the literature and assistance from the community. As an example, for version 3, we relied on PhycoCosm (Grigoriev et al., 2021), MycoCosm (Grigoriev et al., 2014) and PhyloFisher (Tice et al., 2021) to add data sets that became recently available. Because the rate of genome sequencing is increasing exponentially, we will prioritize adding species that are *incertae sedis* and those from taxonomic groups with currently limited representation in EukProt, and only later would include new proteomes from currently well sampled groups.

In the longer term, we hope the standardization of our database provides a path towards including all data sets in a major sequence repository such as NCBI/EBI/DDBJ, so that they can be more broadly accessible and integrated into the suites of tools available at these repositories.

## Acknowledgements

We thank Michelle Leger for helpful advice, Núria Ros-Rocher for suggestions on the figure, Łukasz Sobala and Aleix Obiol for feedback on improvements to the database, Frederick Matsen for helpful discussions, Chris Neely and Harriet Alexander for advice on genome annotation with EukMetaSanity, and members of the Multicellgenome Lab for discussions. We thank the recommender Gavin Douglas and two anonymous reviewers for constructive suggestions that improved this article. Preprint version 5 of this article has been peer-reviewed and recommended by Peer Community In Genomics (https://doi.org/10.24072/pci.genomics.100021).

## Data, scripts and codes availability

The EukProt v3 database is available on FigShare (https://doi.org/10.6084/m9.figshare.12417881.v3). Scripts used for data processing and for building the tree in Figures 1 and 3, as well as annotations of protein data sets (e.g., BUSCO), are available on GitHub via Zenodo (https://doi.org/10.5281/zenodo.7025266). A BLAST server is available at: http://evocellbio.com/eukprot/

